# Bimodality in Ras signaling originates from processivity of the Ras activator SOS without classic kinetic bistability

**DOI:** 10.1101/2023.07.17.549263

**Authors:** Albert A. Lee, Neil H. Kim, Steven Alvarez, He Ren, Joseph B. DeGrandchamp, L. J. Nugent Lew, Jay T. Groves

## Abstract

Ras is a small GTPase that is central to important functional decisions in diverse cell types. An important aspect of Ras signaling is its ability to exhibit bimodal, or switch-like activity. We describe the total reconstitution of a receptor-mediated Ras activation-deactivation reaction catalyzed by SOS and p120-RasGAP on supported lipid membrane microarrays. The results reveal a bimodal Ras activation response, which is not a result of classic kinetic bistability, but is rather driven by the distinct processivity of the Ras activator, SOS. Furthermore, the bimodal response is controlled by the condensation state of the scaffold protein, LAT, to which SOS is recruited. Processivity-driven bimodality leads to stochastic bursts of Ras activation even under strongly deactivating conditions. This behavior contrasts classic kinetic bistability and is distinctly more resistant to pharmacological inhibition.

## Introduction

The small GTPase Ras functions as a molecular switch, toggled between GTP- and GDP-bound states, where only the GTP-bound state is able to interact with and activate downstream effector molecules (*1*). Ras activates a variety of signaling pathways, such as MAPK and PI3K, that affect cellular proliferation, survival, and differentiation (*2, 3*). The activation-deactivation cycle of Ras is dependent on regulatory enzymes that control Ras signaling according to upstream signals (*4, 5*). Guanine nucleotide exchange factors (GEFs), such as SOS, promote the exchange of GDP for GTP to activate Ras (*6, 7*) while hydrolysis of GTP to GDP is promoted by GTPase-activating proteins (GAPs) (*8*). Under physiological conditions, Ras activation and deactivation reactions are in constant competition and must be tightly regulated; misregulation of Ras is a major cause of cancer (*9*). Decades of focused studies on the Ras signaling mechanism have revealed many detailed biochemical insights (*10*), such as its interaction with various effectors (*3, 7, 8*), conformational dynamics (*11, 12*), and its potential to dimerize and form clusters (*13*– *17*). Despite this continuously improving understanding, the Ras signaling mechanism remains enigmatic and difficult to control pharmacologically (*10, 18*).

An important feature of physiological Ras-MAPK signaling is its apparent ability to signal in a bimodal manner (*19*–*24*). This has the effect of converting a continuously variable input to the signal pathway (e.g. the number of activated receptors) into a binary output response and is of extreme importance in general biological information processing (*25*–*27*). In lymphocytes, this bimodal signaling behavior has been ascribed to the feedback kinetics of SOS (*21, 28, 29*) — specifically a RasGTP-driven positive feedback loop (*30*). However, this conjecture is based exclusively on computational modeling and indirect measurements in cellular systems: experimental observation of bimodality in the isolated Ras activation response has not previously been reported (*31*). One contributing factor to the challenges presented by Ras is the surprising complexity that can emerge from competitive enzymatic reaction cycles, especially when a membrane is involved and systems are spatially confined (*28, 32*–*35*).

SOS is a ubiquitously expressed Ras GEF that resides in the cytosol and is recruited to the membrane to initiate Ras activation. SOS is structurally organized with an N-terminal histone-fold (HF), pleckstrin homology (PH) and Dbl homology (DH) domains, a catalytic core with a Ras exchange motif (REM) and a Cdc25 domain, followed by a C-terminal proline-rich (PR) domain (*6*). SOS is subjected to complex regulation, including autoinhibition (*36*) and allosteric activation by Ras (*30*). RasGTP-driven positive feedback in SOS activity has been observed in bulk experiments and attributed to nucleotide selective allosteric activation of SOS by RasGTP (*30*). In earlier modeling studies, SOS was typically assumed to have conventional allosteric activation with different catalytic rates depending on whether RasGDP or RasGTP was bound to its allosteric site (*21, 37*–*39*). When coupled with a competing reaction, this type of positive feedback can lead to kinetic bistability, in which two stable steady states of Ras activity level may exist (*25*). Such a classic kinetic bistability in the activation of Ras by SOS has been presumed to underlie the bimodal signaling response observed at the cellular level in lymphocytes (*21, 28, 29*). The first direct single molecule studies of SOS activity on membranes, however, complicated the story by revealing essentially identical average catalytic rates of SOS with either RasGDP or RasGTP in the allosteric binding site—thus eliminating the possibility of a positive feedback mechanism based on allosteric enhancement of catalytic activity (*37*). Further single-molecule studies with the full-length SOS protein have gone on to resolve a complex autoinhibition release mechanism (*40*), which is coupled to the LAT (*41*) (or EGFR (*42*)) protein condensation phase transition. Once activated, SOS is highly processive and can activate hundreds of Ras molecules during a single membrane binding event (*37, 40, 41*). Collectively, these more recent experimental studies reveal an entirely different activation process in the SOS-Ras system and motivate a comprehensive reevaluation of the mechanistic origins of its bimodal signaling response.

Here, we employ a total reconstitution approach to study the dynamics of Ras activity on a membrane. The experimental system includes the native machinery of receptor-mediated Ras activation by SOS in T cells, consisting of the phosphorylated LAT (pLAT) protein, PIP_2_ lipids, and Ras in a supported membrane along with Grb2, full-length SOS (SOS^FL^) or truncated constructs, p120Gap, and GTP in solution. This system captures the Grb2-mediated recruitment of SOS^FL^ to activated receptor scaffold (pLAT in this case), the membrane-mediated process of SOS autoinhibition release (including PIP_2_ and allosteric Ras binding), SOS-catalyzed nucleotide exchange in Ras, and the dynamic competition with GAP-mediated Ras deactivation. The Ras activity state is read in real-time through the binding of a fluorescently labeled Ras binding domain (RBD) of the downstream effector, Raf (*41*). Experiments are run on supported membrane microarrays, enabling simultaneous measurement of thousands of Ras activation-deactivation reactions at the membrane, all exposed to the same signaling environment from the solution. The microarray strategy also enables observation of stochastic effects in reaction systems of controlled, physiologically relevant sizes.

The experimental results reveal a distinctive scale-dependent bimodality in the Ras activity state with SOS^FL^. The bimodal response is only observed in small corralled reaction systems, indicating that this effect does not originate from classic kinetic bistability (*33*). The system becomes monostable at larger sizes, even while all other parameters are maintained identical. We further observe that the truncated catalytic core of SOS (SOS^Cat^) is incapable of driving a bimodal Ras activity response under any experimentally accessible conditions. SOS^Cat^ includes both the catalytic and allosteric Ras binding sites and exhibits Ras-GTP driven positive feedback, which nonetheless proved insufficient to establish kinetic bistability and is not the primary driver of the observed bimodal Ras signaling response. Through a combination of studies using other SOS truncations, along with stochastic kinetic modeling, we establish that it is the extreme processivity of SOS that enables the bimodal signaling response through a stochastic mechanism. Individual molecular SOS activation events drive bursts of Ras activation, which locally overcome RasGAP activity even under strongly deactivating conditions. This behavior contrasts that of bimodality driven by positive feedback, in which Ras activation is much more distributed. Processivity-driven stochastic bimodality is markedly insensitive to SOS inhibition, suggesting it could present challenges to the development of effective therapeutics to inhibit Ras activation by targeting SOS (*10, 43*–*46*). We also observe that bimodal Ras activity is dependent on system size as well as the local density of Ras or LAT, thus offering more degrees of control than would be available if it were rooted in a classic kinetic bistability. More broadly, these studies reveal how stochastic variation from single molecular SOS activation events can be amplified by the extreme processivity of SOS to drive a full cellular-level bimodal Ras signaling response.

## Results

### Reconstitution of Ras activation-deactivation competition reaction

The activity state of Ras is controlled by competing GEFs and GAPs that primarily interact with Ras on the cell membrane (*4, 5*). Many commonly used Ras biochemical assays utilize solubilized versions of Ras and are carried out as solution assays (*30, 36*). While such assay formats can be convenient, they also eliminate many important regulatory mechanisms (*35, 37, 41*). For example, the ubiquitous Ras GEF, SOS, in its full-length form, is strongly autoinhibited and shows almost no activity in solution (*40, 41*). SOS truncations with impaired autoinhibition must generally be used for observable activity in solution assays. Additionally, once activated, SOS is highly processive and can activate hundreds of Ras molecules during a single membrane binding event (*37, 40, 41*). This processivity is only achievable on membranes and furthermore requires an extended membrane format; conventional vesicles (typically around 100 nm in diameter) can be artificially limiting by allowing SOS to completely activate all available Ras molecules (*47*). Here we seek to reconstitute the entire receptor-mediated recruitment and activation of SOS in a planar membrane format that includes receptor-mediated SOS recruitment via Grb2, lipid- and Ras-dependent autoinhibition release in SOS, as well as the inclusion of competing GAP-driven Ras deactivation.

We reconstituted Ras activation-deactivation competition reaction on a supported lipid bilayer (SLB) containing 92% DOPC, 2% PIP_2_, 2% MCC-DOPE, and 4% Ni–NTA-DOGS (see Materials and Methods for details). H-Ras(C118S, 1-181) (hereafter referred to as “Ras”) is linked to the SLB by coupling the Cys181 to the MCC-DOPE lipids through thiol-maleimide reaction, mimicking the native lipid-modification of Ras (*35*). The resulting membrane-linked Ras is stably bound and fully functional (*35, 37, 41*). The cytoplasmic region of LAT, expressed along with an N-terminal His6 tag, is linked to the SLB through binding between its His tag and Ni-NTA lipid (*48*–*50*). LAT is phosphorylated by a Src-family kinase protein, Hck, which is also tethered to the SLB (*41*). The tethered proteins are laterally mobile, and their surface density on the membrane is calibrated precisely by fluorescence correlation spectroscopy (FCS) (see Materials and Methods for details). Typical densities used in the experiments described here are ∼1500/µm^2^ for Ras and ∼500/µm^2^ for LAT, both of which are comparable to physiological densities found in cells (*35, 51*).

Time evolution of the Ras activation state (here defined as the fraction of activated Ras: RasGTP/(total Ras)) can be measured by quantitative fluorescence imaging of the binding of a Ras binding domain (RBD) sensor to RasGTP using total internal reflection fluorescence (TIRF) microscopy (*41*). Upon the addition of SOS, Grb2, and GTP from the solution, robust Ras activation can be detected (Figure S1A). Kinetic traces of Ras activation by SOS^Cat^ exhibit sigmoidal shapes, confirming that the catalytic core of SOS exhibits a RasGTP-driven positive feedback as reported previously (*30, 36, 42*). In order to examine steady-state Ras activation under competitive activating and deactivating reactions, we introduce the catalytic domain of p120RasGAP (hereafter referred to as “RasGAP”) from the solution. RasGAP drives a simple, essentially bimolecular Ras deactivation reaction, with no evidence of feedback (Figure S1B). The combination of SOS and RasGAP establishes a continuously cycling Ras activation-deactivation competitive reaction (Figure S1C), which is drawn schematically in Figure 1.

**Fig. 1.**
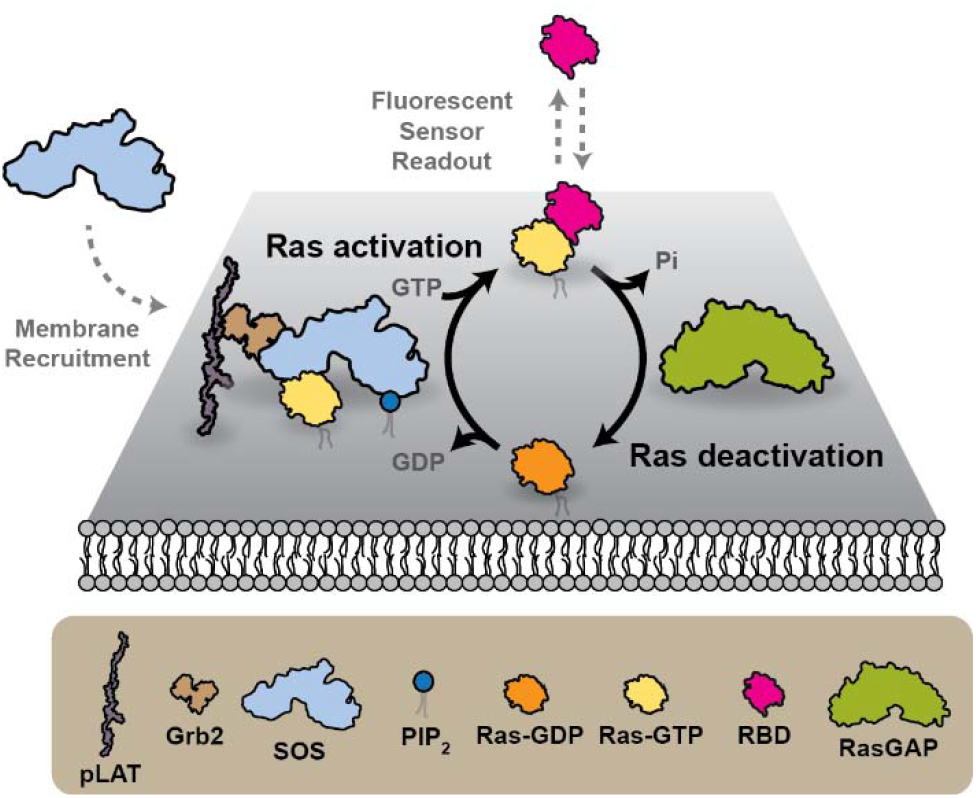
Schematic of the reconstitution of a Ras activation-deactivation reaction. SOS can be recruited to the membrane by Grb2 mediated binding to pLAT and activate Ras. GAP from the solution can directly deactivate Ras. Ras activation can be detected by the binding of fluorescently labeled RBD.

### Ras activity exhibits large fluctuations in microscopically confined reaction systems

Ras signaling reactions under cellular conditions are intrinsically confined on microscopic length scales by the physiological geometry of the cell. Additionally, SOS is estimated to be expressed at only 2000-7000 molecules per cell (*52*). In many cases, such as in T cell activation, signaling reactions are confined to localized signaling clusters on the membrane, further reducing the copy numbers of participating molecules (*53, 54*). Spatial confinement and low molecular copy numbers lead to substantial stochastic variation that effectively changes the laws of chemistry within the cell (*55*), sometimes enhancing sensitivity (*56*), inducing bistability (*28, 57, 58*), or even changing the outcome of a competitive signaling reaction (*33, 34*).

Here we adopt an experimental approach in which system size is a directly controllable parameter by using supported membrane microarrays (*37, 41, 59*) (Figure 2A). In this strategy, a supported membrane is formed on glass substrates prefabricated with arrays of metal lines. Lipids and membrane-bound components, such as Ras, diffuse freely within each confined corral but cannot cross the metal barriers, segregating each corral of membrane into an independent reaction system. Each corral, however, is in contact with an identical bulk solution phase and the metal barriers (only ∼10 nm in height) have no interference with diffusion and flow within the solution phase (*59*). Prior to any reaction process, the initial state of the membrane components exhibits minimal variability across corrals (Figure S2). With the membrane microarray, we are able to simultaneously track thousands of competitive Ras activity reactions, all of which experience essentially identical reaction conditions, with spatial resolution down to ∼1 µm and temporal resolution to 100 ms (Figure 2B).

**Fig. 2.**
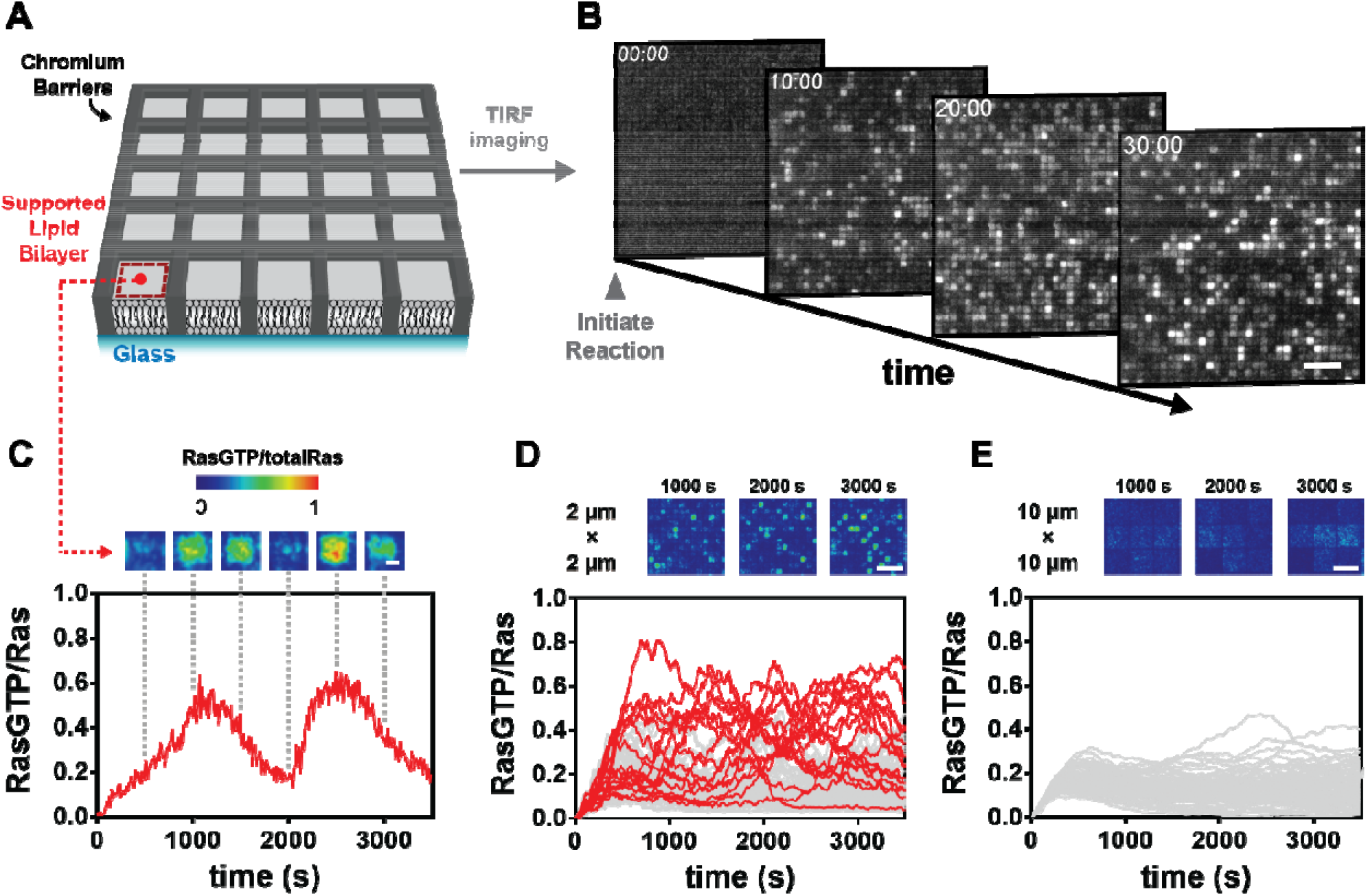
Ras activation-deactivation reaction in membrane microarrays. **(A)** Schematic of the membrane microarray. **(B)** Time course images of Ras activation-deactivation reactions in membrane microarrays. Scale bar 10 µm. **(C)** Ras activation-deactivation reaction in a 2 µm × 2 µm corral. Scale bar 1 µm. **(D)** Reaction trajectories in 2 µm × 2 µm corrals (n = 400). Trajectories that exceeded 0.5 RasGTP/Ras are labeled red. Scale bar 10 µm. **(E)** Reaction trajectories in 10 µm × 10 µm corrals (n = 100). Scale bar 10 µm.

Within smaller corral sizes (e.g. 2 μm × 2 μm), robust stochastic pulses of Ras activation can be observed within individual corrals under steady-state conditions in the competitive activation reaction (Figure 2C). Such fluctuations can drive the Ras activation state far away from the mean and occur on the time scale of hundreds of seconds. An ensemble of 400 steady-state reaction traces from a 2 μm × 2 μm corral array is plotted in Figure 2D. Although the mean Ras activity (RasGTP/(total Ras)) is 0.079, many reaction traces (highlighted in red) cross the threshold of having more Ras-GTP than Ras-GDP (Ras activity = 0.50). This magnitude of stochastic variation cannot result from the intrinsic stochasticity of individual Ras activation reactions. With thousands of Ras molecules in each corral, such intrinsic variation (which scales with the square root of the number of molecules) is limited to a few percent (*60*). Instead, the observed variation is caused by individual SOS molecules entering a highly processive state, where the high catalytic output from a single SOS molecule can locally overwhelm the opposing GAP-catalyzed Ras deactivation reaction. This effect can be directly observed when simultaneously monitoring SOS binding to the membrane (with fluorescently labeled SOS constructs) and local Ras activation (Figure S3). We suggest such temporal spikes in Ras activity in the competitive reaction, here observed to last for hundreds of seconds, could trigger downstream signaling activity even while the average Ras activation level may be insufficient to do so. In larger corrals, different behavior is observed. Reaction traces from an ensemble of 100 10 µm x 10 µm corrals are plotted in Figure 2E. Whereas the mean Ras activity (0.13) is similar to that observed in the 2 µm x 2 µm corrals, we now observe that none of the reaction traces exceed the threshold (Ras activity = 0.5) at any point. In these larger sized corrals, the concentrated activity from individual SOS molecules is effectively diluted out, and the individual reaction traces track much closer to the mean.

### The Ras activation-deactivation reaction exhibits a size-dependent bimodal response

Evidence for bimodal signaling behavior through the Ras-MAPK pathway has been observed in a number of cellular systems (*19*–*24*). It has been proposed that RasGTP-driven positive feedback in SOS could lead to kinetic bistability in a competitive Ras activation-deactivation reaction system, and that this is the origin of cellular bimodal signaling behavior (*21, 28, 29*). However, bimodal Ras signaling in reconstituted systems has not previously been reported. Here we test this hypothesis by directly examining the steady-state Ras activity behavior in competitive SOS-RasGAP reactions under a wide range of conditions and reaction system sizes. Unique features in these experiments that enable complex Ras activity behavior include: ***i***) Grb2-mediated recruitment of SOS^FL^ to activated receptor scaffold (pLAT); ***ii***) fluid movement of SOS on the membrane surface to processively activate many Ras molecules; and ***iii***) control of reaction size with the micropatterned supported membrane system.

We initially investigate Ras activation-deactivation reactions in membrane microarrays ranging from 1 μm × 1 μm up to 10 μm × 10 μm. The reaction systems were allowed to reach an apparent steady-state (after ∼2 hr), at which point we examine the spectrum of corral-to-corral variability of the Ras activity state (Figure 3A). These reactions are run under conditions where the average Ras activity in macroscopic reactions is roughly 0.5 (upper images in Figure 3A). Probability distributions for the Ras activity state, derived from histograms of the observed activity on the corral arrays, are plotted below each image in Figure 3A. We observe the steady-state Ras activity to be bimodally distributed in the 1 μm × 1 μm and 2 μm × 2 μm corral arrays while becoming unimodal in larger arrays. This scale dependence of the observed bimodality is not consistent with classic kinetic bistability. If there were a stable kinetic bistability underlying the bimodal signaling behavior, this would still be observed in the larger size corrals—appearing as spontaneously separating domains of Ras activity state (see, for example, the PIP_1_/PIP_2_ lipid patterns generated from kinetic bistability described in (*33*)). Instead, the observation of robust bimodality in small corrals and unimodality in larger corrals indicates a stochastic origin to the bimodal behavior in a system that is intrinsically monostable (*33, 34*).

**Fig. 3.**
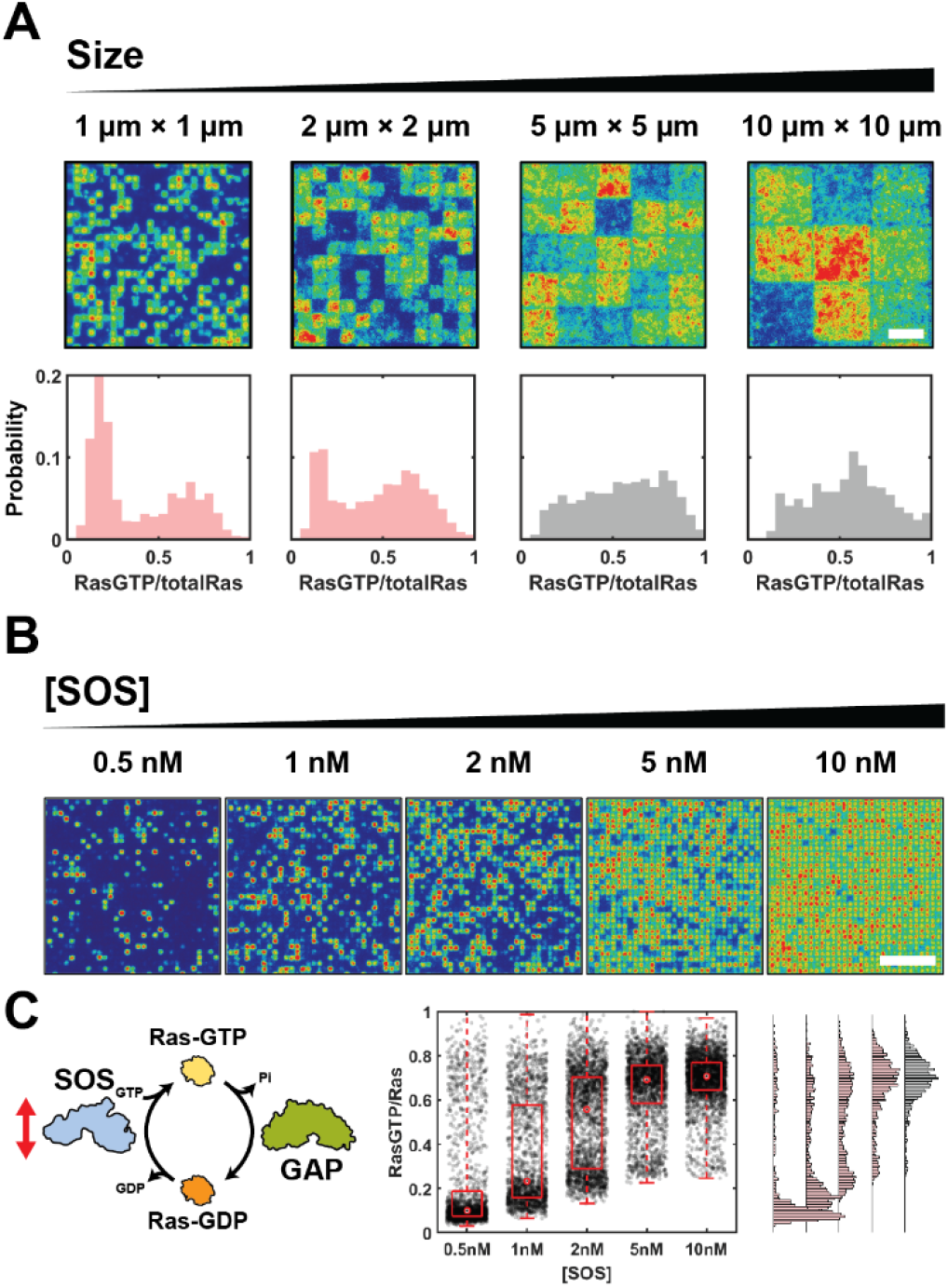
Bimodal Ras activation in the Ras activation-deactivation reaction. **(A)** Distribution of Ras activation states in the Ras activation-deactivation reaction in corrals (n > 216). Conditions that are bimodal (based on the Hartigans’ dip test (p < 0.01)) are plotted in red. Scale bar 5 µm. **(B)** Images of the Ras activation-deactivation reaction in 1 μm × 1 μm corrals with varying SOS concentration. Scale bar 10 µm. **(C)** Distribution of Ras activation states in Figure 3B. Each dot represents a single corral. Right: Histogram of the Ras activation state distribution. Conditions that are bimodal are plotted in red.

We next examine the robustness of the bimodal Ras activity response over a range of SOS concentrations, focusing on the 1 μm × 1 μm corral size. At the lowest solution concentration of SOS we examined (0.5 nM), the majority of the reactions have essentially no Ras activation, but a rare subset with distinctly high Ras activation states are clearly resolved (Figure 3B). As the SOS solution concentration is titrated from 0.5 nM up to 10 nM, we observed a change in the ratio of low and high Ras activity states (Figure 3B). The data for this titration of SOS against a fixed RasGAP concentration in a competitive reaction at steady state are summarized in Figure 3C. The system exhibits a bimodal response over the entire range of SOS concentrations. The high Ras activity state is at essentially the same activity level under all conditions but is observed with higher frequency at higher SOS levels. The lower Ras activity state shifts to higher activity levels (but still firmly resolved from the high activity state) and lower frequency with increasing SOS concentration.

### The processivity of SOS is central to the bimodal response

We hypothesize that the observed bimodality in Ras activation is driven by the extreme processivity of SOS. This hypothesis is partly motivated by the fact that bimodality is only observed in smaller reaction systems, which is suggestive of an underlying stochastic mechanism (*33, 34*). SOS processivity is an intrinsic amplifier of stochasticity as it translates single-molecular SOS activation events into bursts of hundreds of activated Ras molecules. We test this hypothesis by examining three truncated SOS constructs (Figure 4A): ***i***) SOS^Cat^, only the catalytic domains with both the N- and C-terminal regulatory domains truncated; ***ii***) SOS^CatPR^, with the N-terminal regulatory domains truncated; and ***iii***) SOS^HDPC^, with the C-terminal regulatory domains truncated (Figure 4A). All constructs, including SOS^Cat^, have both allosteric and catalytic Ras binding sites and can exhibit RasGTP-driven positive feedback. The inclusion of the PR (SOS^CatPR^) or the N-terminal regulatory domains (SOS^HDPC^) domains adds autoinhibition as well as additional membrane binding interactions, which enhance SOS processivity. The C-terminal PR domain allows SOS^CatPR^ to bind to the membrane through Grb2 and phosphorylated LAT. The N-terminal HF, PH, and DH domains allow SOS^HDPC^ to bind to the membrane through lipids such as PIP_2_.

**Fig. 4.**
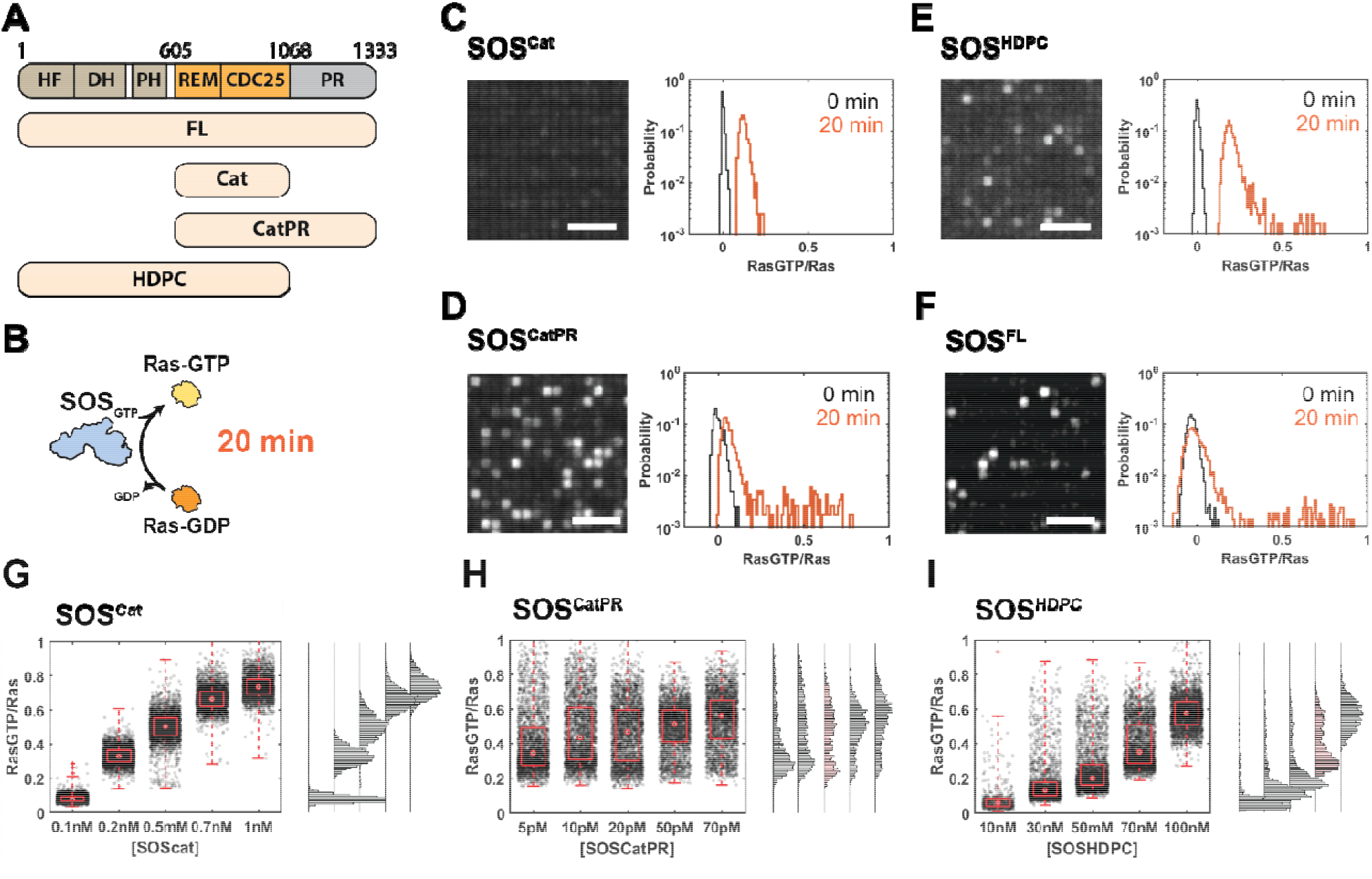
Reactions of truncations of SOS with varying processivity. **(A)** Truncations of SOS. **(B)** Time limited one-way reaction in Figure 4C∼4F. **(C)** SOS^Cat^ reaction. Left: Image of the reaction at 20 min. Right: Distribution of Ras activation states. The black histogram is the distribution at 0 min. The orange histogram is the distribution at 20 min. Scale bar 5 µm. **(D)** SOS^CatPR^ reaction. **(E)** SOS^HDPC^ reaction. **(F)** SOS^FL^ reaction. **(G)** Distribution of Ras activation states in SOS^cat^-p120GAP competition reaction. **(G)** SOS^CatPR^-p120GAP competition reaction. **(G)** SOS^HDPC^-p120GAP competition reaction. Histograms that are bimodal are plotted in red.

The SOS constructs exhibit different levels of processive activity due to the different types of membrane association, with SOS^FL^ exhibiting the highest processivity (*40, 61*). The truncated SOS constructs can also activate Ras in a non-processive manner, through transient collisions with the membrane. This activity is most prominent in SOS^Cat^, which lacks all autoinhibition, and is progressively lower for both SOS^CatPR^ and SOS^HDPC^, each of which has some autoinhibition. Transient activity is essentially unmeasurable in SOS^FL^, due to its thorough autoinhibition. These different activity profiles for the SOS constructs can be resolved in an assay where SOS is allowed to catalyze Ras activation, in a one-way reaction without competing RasGAP, for a defined time (20 minutes) in membrane microarrays (Figure 4B). Results comparing initial and final Ras activity states for the four SOS constructs in 1µm x 1µm microarrays are illustrated in Figures 4C-F. The distribution of the Ras activation states sampled from thousands of corrals is plotted to the right of each representative image of the corral array. In the case of SOS^Cat^ (Figure 4C), the Ras activity levels exhibit a Gaussian distribution centered around a well-defined mean after 20 minutes of activation. There are no outlier high Ras activity corrals for SOS^Cat^. This behavior corresponds with primarily transient activity, which uniformly distributes the Ras activation events among the corrals. For SOS^CatPR^ (Figure 4D) and SOS^HDPC^ (Figure 4E), there is a primarily Gaussian peak around a moderate Ras activity level (from transient activity) with a number of high Ras activity corrals far outside this distribution, which correspond to occasional SOS molecules entering the processive state. SOS^FL^ exhibits the most extreme behavior, with a Gaussian peak centered at essentially zero Ras activity and a number of highly activated corrals from processive SOS molecules (Figure 4F). These data confirm that the processive state is the only active state in native, SOS^FL^ (*40, 41, 62*).

In competitive reactions, we observe the processive activity of SOS is instrumental in overcoming the deactivating pressure from RasGAP to achieve high Ras activity states. Titrating SOS^Cat^ solution concentration in the presence of RasGAP on 1µm x 1µm corral arrays (as done previously with SOS^FL^, see Figure 3) reveals a continuously shifting Ras activity level, with no evidence of bimodality at any SOS^Cat^ concentration (Figure 4G). Similar titrations with SOS^CatPR^ and SOS^HDPC^ reveal weak bimodality in some conditions (Figure 4H and 4I). Only SOS^FL^, which exclusively activates Ras through the processive state, exhibits clear bimodality over the entire range of concentrations sampled (see Figure 3C).

### Stochastic simulations confirm bimodality results from processivity

To further examine the connections between processivity and bimodality in Ras activation by SOS, we conducted a series of stochastic simulations. We used a simplified model that includes only the common processes among all SOS constructs. In this model (Figure 5A), SOS in solution (SOS_sol_) can reversibly bind to either RasGDP or RasGTP through its allosteric site to become localized at the membrane; these membrane-associated species are labeled SOS:RasGDP and SOS:RasGTP, respectively. The binding affinity to the SOS allosteric site of RasGTP is stronger than RasGDP (*30, 36*) and this is reflected in the model. SOS:RasGDP or SOS:RasGTP catalyze the conversion of RasGDP to RasGTP with the same apparent rate constant, 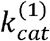, which has been experimentally measured in single-molecule membrane array experiments (*37*). Although 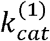 is independent of the allosteric Ras nucleotide state, and therefore can make no contribution to positive feedback, this model still exhibits RasGTP-driven positive feedback through the differential binding of RasGTP vs. RasGDP at the SOS allosteric site and the fact that an allosterically bound Ras (of any nucleotide state) is required for catalytic activity (*63*). RasGAP catalyzes RasGTP to RasGDP reaction with simple bimolecular kinetics at a constant rate, 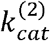, which is experimentally determined from data shown in Figure S1B. Simulations were conducted using the Gillespie algorithm and kinetic rate constants used in the simulation were derived from experimentally determined values (see Materials and Methods for details). We first performed the stochastic simulation with slow 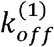 and 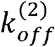, reflecting the experimental measurements, which lead to processive Ras activation by SOS. Under this condition, the stochastic simulation is clearly reflective of the experimental results (Figure 5B), in which the Ras activation state is bimodally distributed only in small reaction systems. By contrast, when the same kinetic parameters and reaction scheme are modeled with deterministic rate equations and concentrations are treated as continuous variables, the system has one unique steady-state solution—kinetic bistability is not achieved. Similarly, when the size of the reaction increases in the stochastic simulation, the Ras activation states become unimodally distributed around the deterministic solution.

**Fig. 5.**
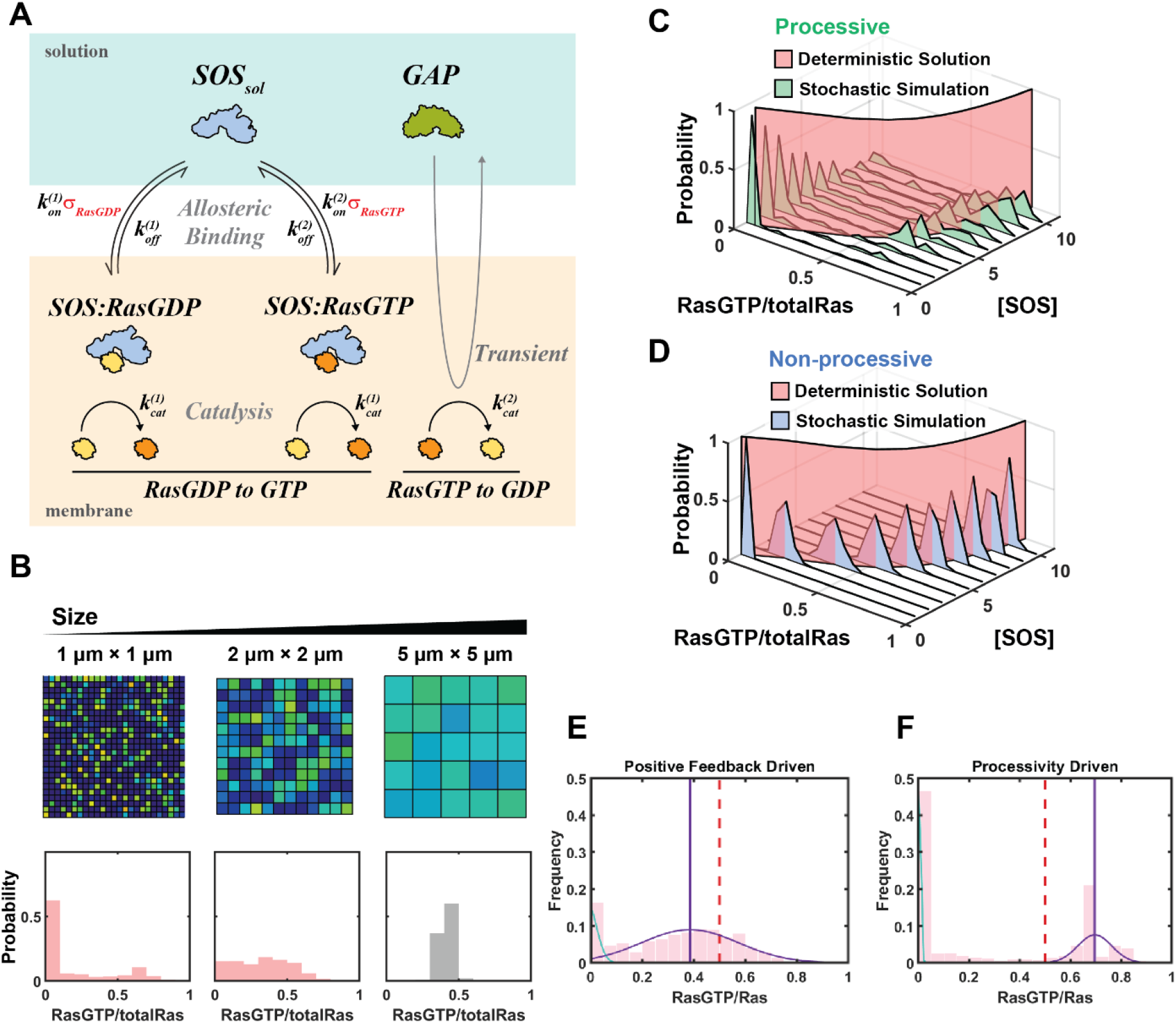
Stochastic simulations of Ras activation-deactivation reactions. **(A)** Kinetic scheme for the stochastic kinetic modeling. **(B)** Simulated result of Ras activation-deactivation reaction in corrals (n>500). Conditions that are bimodal are plotted in red. **(C)** Simulated result of Ras activation-deactivation reaction with varying concentration of processive SOS and **(D)** non-processive SOS in 1 μm × 1 μm corrals. The red plane corresponds to the deterministic steady-state solution. **(E)** Simulation of Ras activation-deactivation reaction with weakly processive SOS that has strong positive feedback and **(F)** with processive SOS that has no positive feedback. The histogram is fitted to two gaussian distribution. The purple line shows the peak of the high mode. The red dashed line shows the deterministic steady-state solution.

We can use the stochastic simulation to isolate the effect of SOS processivity by increasing both *k*_*off*_ and *k*_*on*_ for Ras binding to the allosteric site, while preserving their ratio (*K*_*D*_). Under these conditions, SOS dwells at the membrane so briefly that on average less than one Ras molecule is activated during an individual membrane binding event (no processivity). However, by preserving *K*_*D*_, the total amount of membrane recruited SOS and the overall rate of Ras activation is maintained. Modeling the system with deterministic rate equations yields identical results, irrespective of the degree of processivity of SOS. However, stochastic simulations yield drastically different behaviors. With processive SOS, the Ras activation state distribution responds to an increase in SOS concentration in a bimodal manner (Figure 5C). Conversely, when SOS is non-processive, the Ras activation state distribution responds to an increase in SOS concentration in a unimodal and graded manner that centers around the deterministic solution (Figure 5D). The non-processive SOS behavior closely matches the experimental results for SOS^Cat^ (Figure 3B).

Overall, these simulation results confirm SOS processivity is a key driver of the observed bimodal signaling response and that the system does not exhibit classic kinetic bistability. Broadly speaking, this type of behavior has been termed stochastic bistability (*57, 58, 64*), referring to the fact that bimodal behavior can emerge as a result of stochastic effects in systems that lack intrinsic kinetic bistability according to deterministic rate equations. However, there are different mechanisms by which stochastic effects can lead to bimodality.

One well-known stochastic bistability mechanism is driven by positive feedback (*58*). In this scenario, one of the states will be close to 0 while the other state is near the deterministic steady-state solution. This can be readily reproduced by simulating the reaction in Figure 5A with strong positive feedback and weak processivity (*K*_*D*_ for RasGDP for *K*_*D*_ RasGTP, *K*_*off*_ > *k*_*cat*_. See Materials and Methods for details) (Figure 5E). Compared with the deterministic solution, positive feedback-based stochastic bistability will stochastically deactivate the reaction, populating the 0 state.

Another type of stochastic bistability is driven by processivity (equivalent to burst amplitude in gene expression) (*64*). In the case of processivity-based stochastic bistability (simulating with for RasGDP = *K*_*D*_ for RasGTP, *K*_*off*_ < *k*_*cat*_; see Materials and Methods for details) the deterministic solution lies between the two modes (Figure 5F). Our experimental observations that the Ras activation state in small reaction sizes bifurcates into two modes with one higher and one lower than the Ras activation state in large reaction sizes is consistent with this later mechanism of stochastic bistability, driven by enzymatic processivity.

The subtle differences between these two mechanisms of stochastic bistability can become paramount in a cellular signaling context. Inhibition of SOS is of significant interest as a potential treatment of Ras signaling-related diseases (*10, 43*–*46*). We simulated the effect of SOS inhibition on Ras signaling under conditions of either a positive feedback-based or a processivity-based stochastic bistability (matching conditions in figure 5E and 5F with the addition of reversible inhibition of SOS catalysis, see Materials and Methods for details). In the case of positive feedback-based stochastic bistability, both Ras activity modes are equal or lower than the deterministic solution and the Ras activity of the overall population becomes inhibited relatively easily, as predicted by the deterministic solution (Figure S4A). However, in processivity-based stochastic bistability, even at a drug concentration where deterministic kinetics predict successful lowering of Ras activation, Ras can still be stochastically activated in sufficient small reaction systems (Figure S4B). Over time, essentially every Ras reaction subsystem will experience locally high activation levels, even while the overall mean of the population is highly inhibited. This may provide a hint as to why effective SOS inhibition strategies to control Ras activity have been challenging to develop (*10, 46*).

### Control points for bimodal Ras activation

The bimodal Ras activation response is dependent on Ras density (Figure 6A). The observations of bimodal Ras activity described throughout this study occurred at relatively high Ras densities of 1500∼2000/µm^2^. Such high densities of Ras have been reportedly observed in Ras clusters in cells (*13, 17*), but are distinctly higher than the average Ras density in the plasma membrane (∼400 molecules/µm^2^) (*21*). At a Ras density of 400 molecules/µm^2^, we observe steady-state Ras activity under a competitive SOS^FL^-RasGAP reaction is no longer bimodal and exhibits a graded response with changing SOS^FL^ solution concentration (Figure 6B). When the density of Ras is increased to ∼1800 molecules/μm^2^, the bimodal response is restored. These data indicate that increasing the local Ras density can alter the Ras signaling behavior from graded to switch-like. We speculate that this is due to the elevated encounter rate of SOS and Ras at high Ras density, which effectively speeds up the kinetic transition to SOS processive activity (Figure 6A).

**Fig. 6.**
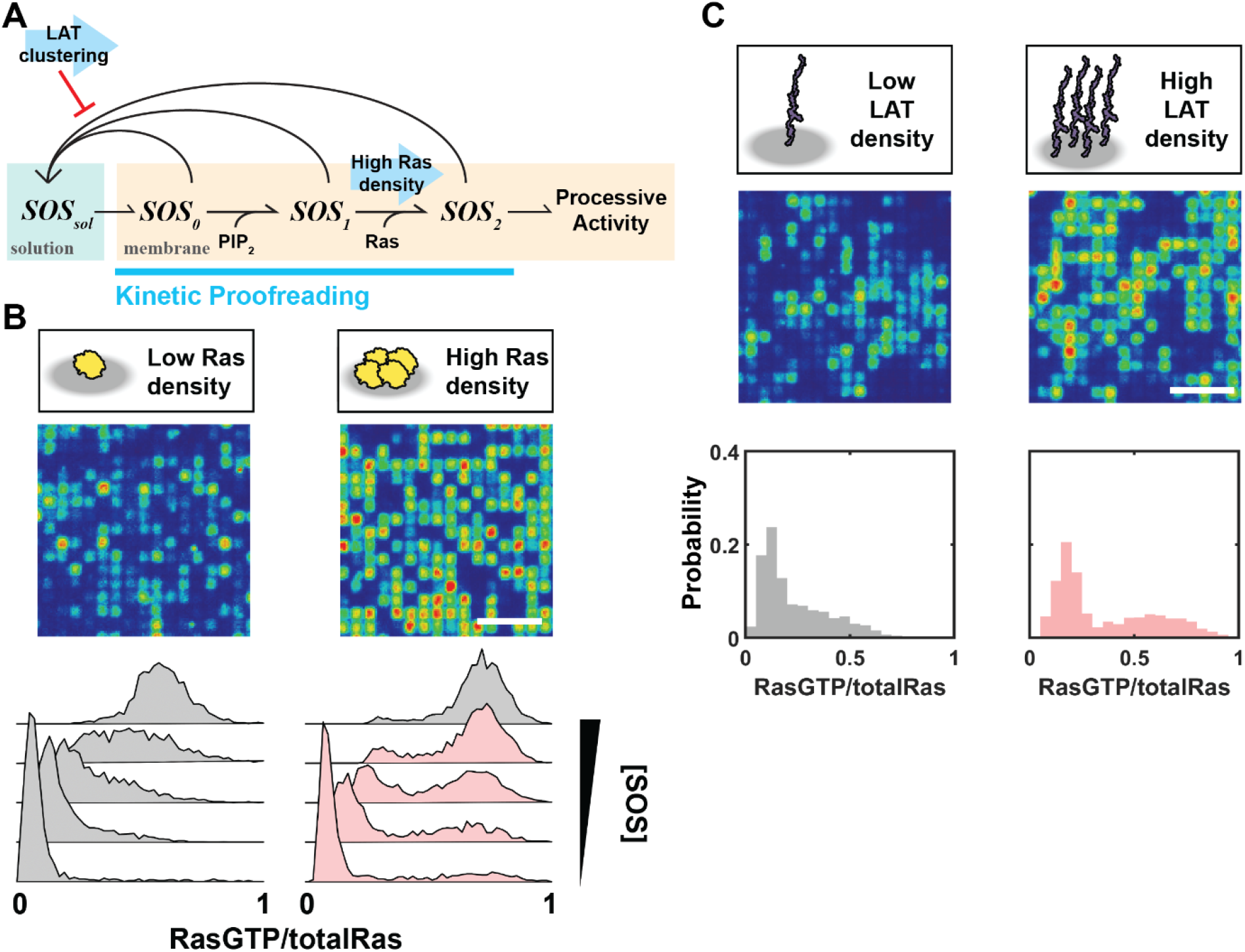
Control points for bimodal Ras activation. **(A)** Kinetic model for SOS activation. SOS needs to bind to the membrane, release autoinhibition, then bind to Ras at its allosteric site before it can become fully activated. These processes are competed by unbinding of SOS from the membrane. **(B)** Ras activation-deactivation reaction with low (∼400 μm^2^) and high (∼2000 μm^2^) Ras density. Scale bar 5 μm. **(C)** Ras activation-deactivation reaction with low (∼500 μm^2^) and high (∼2500 μm^2^) LAT density. Scale bar 5 μm.

Bimodality in Ras activation is also controlled by LAT density (Figure 6A). The scaffold protein LAT forms a two-dimensional protein condensate with Grb2 and SOS on the membrane surface under the control of T cell receptor activation (*41, 49, 53, 65*). A similar condensate has recently been reported with EGFR (*42*). Both the LAT and EGFR condensates control the ability of SOS to activate Ras, through a kinetic proofreading mechanism that taps into the slow autoinhibition release process in SOS. LAT (or EGFR) are effectively clustered in the condensed state and this facilitates multivalent engagement of SOS, which retains SOS at the membrane longer and facilitates autoinhibition release (*41*). We examined the effects of LAT clustering on membranes with low Ras density (∼400 molecules/μm^2^), such that the reaction does not show bimodality under lower, dispersed LAT density (∼500 molecules/μm^2^) (Figure 6C). However, at a higher LAT density (∼2500 molecules/μm^2^, comparable to the density of LAT in the LAT-Grb2-SOS protein condensate (*49*)), the Ras activity state exhibits a bimodal response, despite the low Ras density. In this case, we speculate that the increased activation of SOS into its processive state by the high density of LAT is responsible for the observed bimodality.

Collectively, these results reveal that molecular clustering of LAT or Ras at the membrane can activate the switch-like response from bimodal Ras signaling. We note that this is only possible since the bimodality of Ras activity originates from a stochastic mechanism. If instead it had originated from classic kinetic bistability, the bimodality and switch-like activity would be robust and not sensitive to such subtle perturbations (Figure S5).

## Discussion

Classic kinetic bistability results when nonlinearities in the reaction rates in a competitive reaction cycle produce two, stable steady states. Positive feedback in enzymatic rate is a simple way of generating kinetic bistability (*25*). This phenomenon is widespread in biological signaling systems, where it provides an underlying physical basis for switch-like activity profiles. We have identified an alternative, intrinsically stochastic, mechanism that establishes a bimodal, switch-like, activity response in Ras. This mechanism is dependent on the highly processive activity of SOS and is not the result of RasGTP-driven positive feedback and kinetic bistability. The low copy number of SOS (*52*) and the microscopic reaction scale in cells suggest such stochastic mechanisms likely dominate physiological Ras activity behaviors. This processivity-driven stochastic bimodality achieves similar macroscopic behavior as kinetic bistability, but through a different physical mechanism and with different response characteristics.

A unique feature of the stochastic bistability mechanism is that this allows additional avenues for the switch-like response of Ras signaling to be regulated. We have demonstrated both Ras clustering and LAT clustering, as well as overall reaction size, can serve as control points to engage or disengage the bimodal signaling response. This is distinct from classic kinetic bistability, with two stable steady states, where the switch-like response will always be present. While deterministic kinetic bistability is more robust over a wide range of conditions, stochastic bistability may be more accessible to regulatory control.

A deeper examination of the origin of stochastic bimodality mechanisms clarifies the difference between two types of driving forces: positive feedback and processivity. Stochastic bimodality driven by positive feedback will stochastically deactivate compared to a macroscopic reaction. On the other hand, in SOS-mediated Ras signaling, the stochastic bimodality is driven by processivity and can stochastically activate Ras signaling compared to the macroscopic reaction. This feature may allow Ras signaling to be activated even under a globally inhibiting background of high GAP activity. However, by the same logic, this mechanism may also result in resistance to attempts to inhibit Ras with drugs.

Despite numerous simulation and theoretical studies on stochastic bimodality (*57, 58, 66*), there have been very few experimental reports that describe stochastic bimodality in biological signaling systems (*64, 67*). In part, this is due to the difficulties in analyzing biological reactions in a controlled microscopic environment where physiologically low molecular copy numbers are preserved. The present study describes a systematic approach to studying stochastic effects in membrane signaling reactions based on membrane microarrays. Many signaling proteins exhibit complex regulation at the membrane, including phospholipase C-γ (*68*), phosphoinositide 3-kinases (*69*), and N-WASP (*70*). We speculate that many of the complex molecular features of these signaling proteins could induce bimodal responses in a cellular reaction environment even when classic kinetic bistability can be ruled out.

## Materials and Methods

### Protein purification

SOS and SOS^CatPR^ (533–1333) were purified based on protocols described in previous work (*35*). H-Ras (1-181, C118S) (human H-Ras protein with residues 1-181 and a point mutation to serine at residue C118), SOS^Cat^ (566-1049), SOS^HDPC^ (1-1049) were purified based on previous reports (*35*). The GAP domain of p120GAP (714-1047) was expressed and purified based on the protocols described in previous work (*8*). RBD (56-131; K65E) derived from the Raf-1 human gene was purified and labeled with Alexa647-maleimide using previously reported methods (*41*). LAT (30-233) and Grb2 were purified and labeled following previously described protocols (*41, 69*). Hck was purified based on published protocols (*71*).

### Preparation of liposome

1,2-dioleoyl-sn-glycero-3-phosphocholine (DOPC), L-α-phosphatidylinositol-4,5-bisphosphate (Brain PI(4,5)P_2_), 1,2-dioleoyl-sn-glycero-3-[(N-(5-amino-1-carboxypentyl)iminodiacetic acid)succinyl] (nickel salt) (Ni-DGS), 1,2-dioleoyl-sn-glycero-3-phosphoethanolamine-N-[4-(pmaleimidomethyl)cyclohexanecarboxamide] (MCC-PE) were purchased from Avanti Polar Lipids. Lipids were mixed in a glass round bottom flask cleaned by piranha etching at the desired molar fraction. The solution was then evaporated using a rotary evaporator for 10 min at 35°C or until dried to a thin film. Dried lipid films were further blow-dried with N_2_ for at least 10 min. Lipids were resuspended in Milli-Q H_2_O by shaking and gently pipetting to form a solution with a final concentration of 1 mM total lipids. Small unilamellar vesicles (SUV) for the formation of supported lipid bilayer were prepared by sonication for 100 sec (20 sec on, 30sec off for 5 times) in an ice-water bath.

### Microfabrication

Chromium patterns (100 nm thick and 5 nm high) were fabricated onto 25×75 mm glass coverslips by the Pulsed Nanoimprint Lithography method (Pulsed NIL) (ThunderNIL Srl, Italy). Briefly, a stamp with desired patterns was fabricated by electron beam lithography and was treated with hydrophobic trichlorosilane to make it non-adhesive. Pulsed NIL was performed on glass substrates, which were previously spin-coated with 120 nm thick mr-l 7010 resist, using the stamp. Residual resist film on the glass substrate was etched off using oxygen plasma before the chromium lift-off process.

### Supported lipid bilayer experiments

Glass coverslips with chromium patterns were etched with piranha solution for 5 minutes and then rinsed with water extensively. The coverslip was blow-dried with nitrogen gas and immediately attached to a flow chamber (Ibidi µ-Slide 80608). SLBs were formed on a glass substrate by flowing around 150 μL of 0.25 mM SUVs diluted in PBS (pH 7.2) into the chamber and incubated for at least 30 min. After incubation, the chambers were washed with 1 mL of PBS and then blocked with 1 mg/mL β-casein (Thermo Fisher Scientific 37528) for 10 min. The chambers were then rinsed with 1 mL PBS buffer. H-Ras was incubated at 0.1∼1 mg/mL for 2 hr 30 min in PBS buffer at room temperature. The chambers were then washed with 1 mL PBS, and 5 mM BME was subsequently added to quench the reaction. After 10 min, the flow channel was washed with 1 mL of PBS, and buffer exchanged into the reaction buffer (40 mM HEPES (pH 7.4), 100 mM NaCl, 5 mM MgCl_2_) plus 100 μM GDP. 10∼200 nM LAT and 12.5 nM Hck are introduced into the flow channel and incubated for 40 min. After washing with 1 mL reaction buffer, the flow channels were incubated with 1 mM ATP and 100 μM GDP to phosphorylate LAT and ensure nucleotide loading in Ras. ATP and GDP were washed away with the reaction buffer plus 10 mM BME and immediately before the reaction. All Ras reactions were performed in a buffer containing 40 mM HEPES (pH 7.4), 100 mM NaCl, 5 mM MgCl_2_, 10 mM BME, 100 μM GTP, 2 mM UV-treated Trolox, with the addition of 20 nM Alexa647-RBD(K65E) to monitor the reaction. For SOS and SOS^CatPR^ reactions, 20nM Grb2 was included as well. In competition reaction with p120GAP, 200 nM p120GAP was used.

### Microscope hardware and imaging acquisition

TIRF imaging experiments were performed on an inverted Nikon Eclipse Ti microscope using either a 100x Nikon objective (1.49 NA) oil immersion TIRF objective or a 60x Apo TIRF oil immersion objective (1.45 NA). The light sources were either a 488 nm, 561 nm, or 637 nm diode laser (OBIS laser diode, Coherent, Santa Clara, CA) controlled with a custom-built Solamere (Salt Lake City, Utah) laser driver with analog and digital modulation (0-5 volts). Images were acquired on an EMCCD camera (Andor Technology Ltd., UK). All microscope hardware was controlled using Micro-Manager v4.0. Samples were excited with 0.3∼0.8 mW laser power at the objective. The exposure time is typically 100∼300 ms. In end point assays, imaging starts after 2hrs of incubation to avoid photobleaching.

### Surface Density Measurements

The surface density of Ras and LAT were measured using fluorescence correlation spectroscopy (FCS) on a homebuilt setup based on a Nikon Eclipse TE2000-E inverted microscope. A pulsed white light laser source (SuperK Extreme EXW-12, NKT Photonics) was filtered by bandpass filters for desired excitation wavelengths and combined through a single-mode optical fiber. The excitation pulses enter the microscope via a multicolor dichroic cube (Di01-R405/488/561/635-25×36, Semrock). The fluorescence signal is collected by a 100x high-numerical aperture oil-immersion objective (Nikon), recorded by avalanche photodiode detectors (Hamamatsu), and directly converted into an autocorrelation signal by a hardware correlator (Correlator.com). The resulting autocorrelation G(t) was fit to the two-dimensional Gaussian diffusion model to calculate surface density. LAT densities were measured by directly labeling LAT with Alexa-555 dye. Ras densities were measured by loading Ras with Atto-488-GDP (Jena Bioscience) using SOS^Cat^. Surface density calibration was achieved by fitting the FCS surface density measurements to the TIRF intensity measurements with a linear regression.

### Stochastic Simulation

The time evolution of all species in the reaction was simulated stochastically using the Gillespie algorithm in MATLAB, following the kinetic scheme in Figure 5A. Two assumptions were made: 1) The membrane compositions are spatially homogeneous within reaction space due diffusion. 2) the solution concentration of *SOS*_*sol*_, 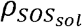, is constant since there is a large solution reservoir in the experiment. Each molecular species is expressed as the exact number of molecules. The rate for each transition is calculated as follows:

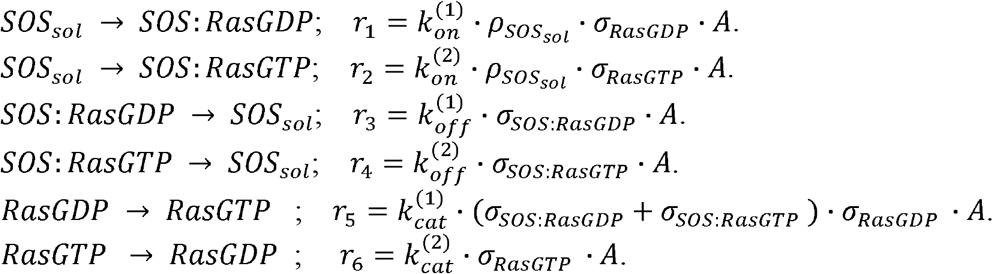

*A* is the area of the membrane in μm^2^, and the surface density of each membrane-associated species, *σ*_*x*_, is expressed as a discrete molecular copy number per unit area. We used the following rate parameters:

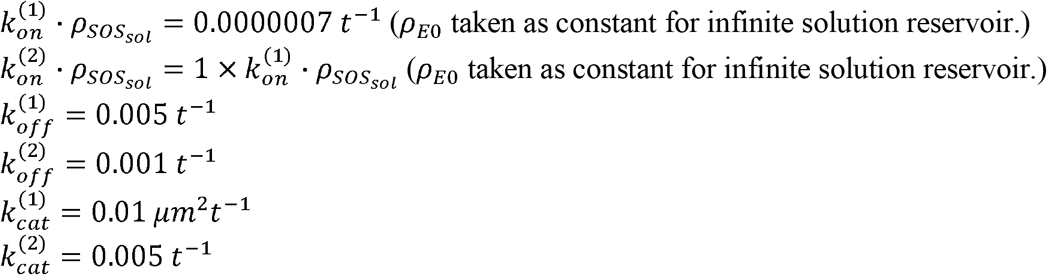

The kinetic parameters used reflect reported kinetic rate constants (*35, 37, 61*). All simulations begin with 1000/μm^2^ RasGDP. Statistics were collected from 500 simulations. In concentration titration simulations, 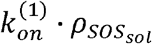 varies from 0.0000003∼0.00003 t^-l^.

For simulations with non-processive SOS, we used the following rate parameters:

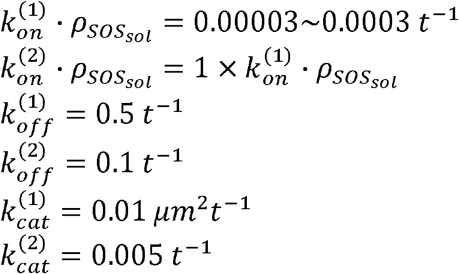

For the simulations of Fig 5E and 5F, we modified the simulation parameters to facilitate comparison. Specifically, we used an adjusted 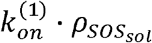 that leads to 0.5 RasGTP/Ras steady state response in deterministic kinetics. We also make the positive feedback strictly from the difference in 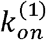 and 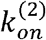, and leaving 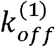 and 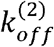 the same to eliminate the dependence of processivity on changes in feedback strength.

For Fig 5E, the rates used are:

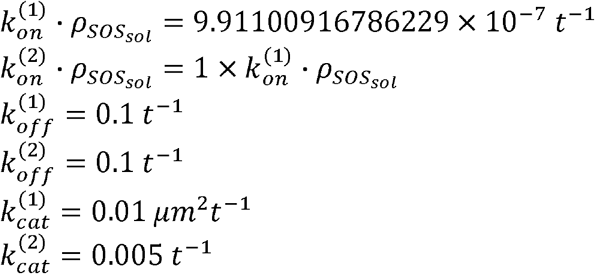

For Fig 5F, the rates used are:

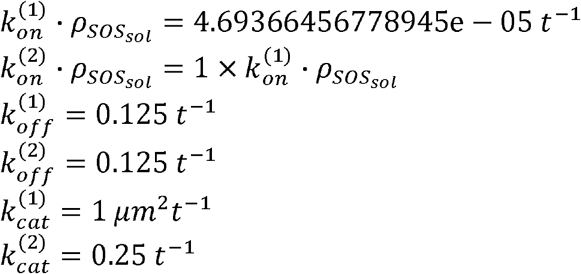

For the simulation in Figure S4A with no inhibitor, we used the following rate parameters:

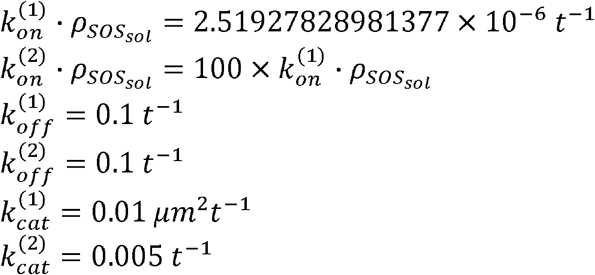

For the simulation in Figure S4B with no inhibitor, we used the following rate parameters:

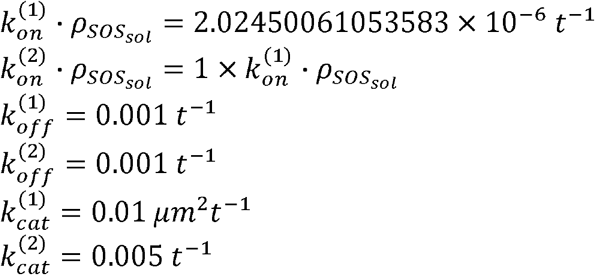

For the simulation of Ras activation-deactivation reaction with SOS inhibitor, the following transitions are simulated:

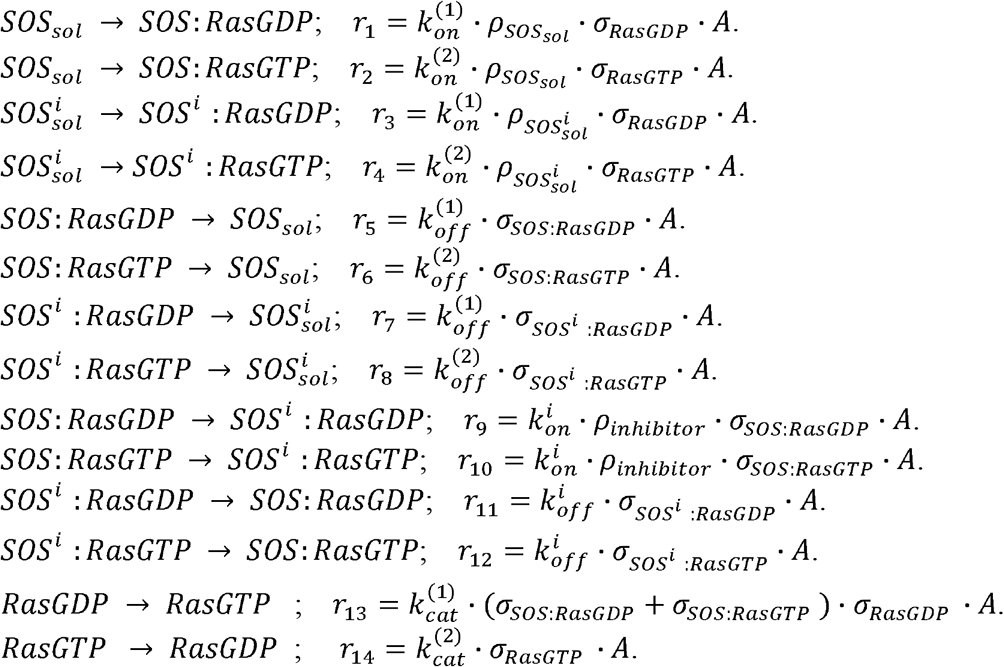

SOS species with superscript “i” are bound to the inhibitor and are catalytically inactive. The solution concentration of the inhibitor, *ρinhibitor* and the solution concentration of 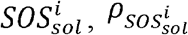, are constant. This is assuming the inhibitor binding has reached equilibrium in the large solution reservoir. For the simulation in Figure S4A with no inhibitor, we used the following parameters:

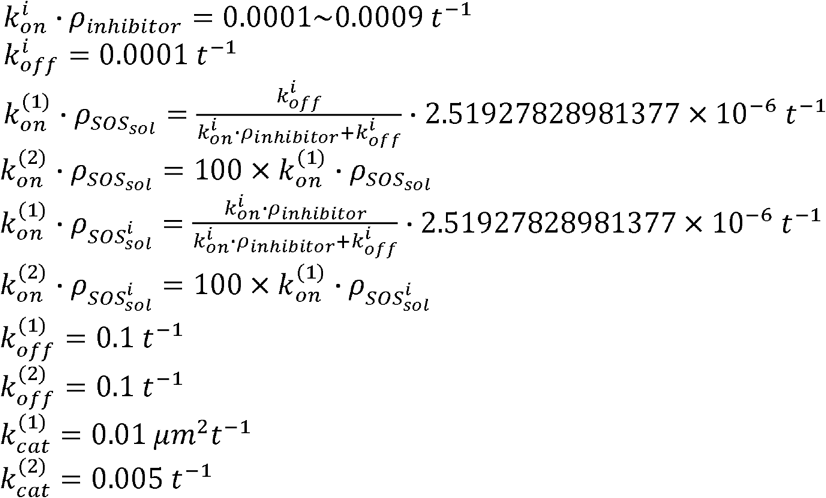

### Deterministic Steady State Solution

The deterministic steady state solution is obtained by solving the following equations using MATLAB.

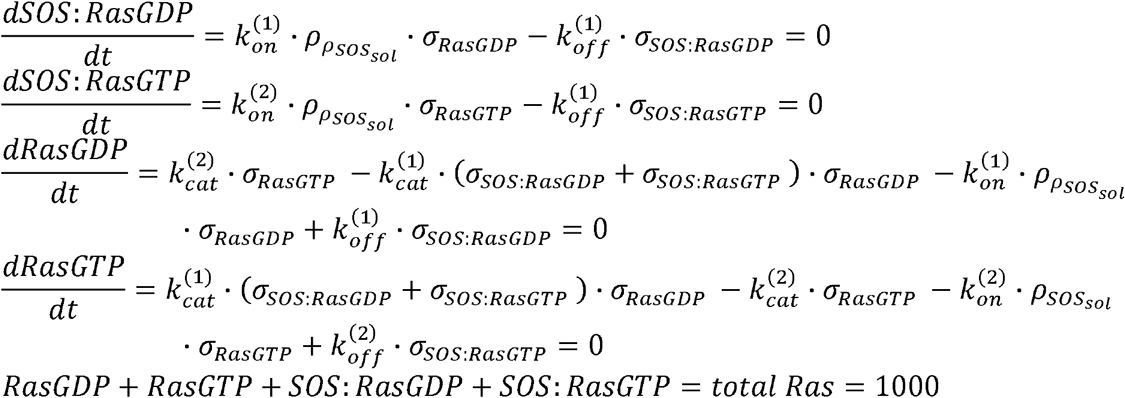

For figure S5, the equations are modified to:

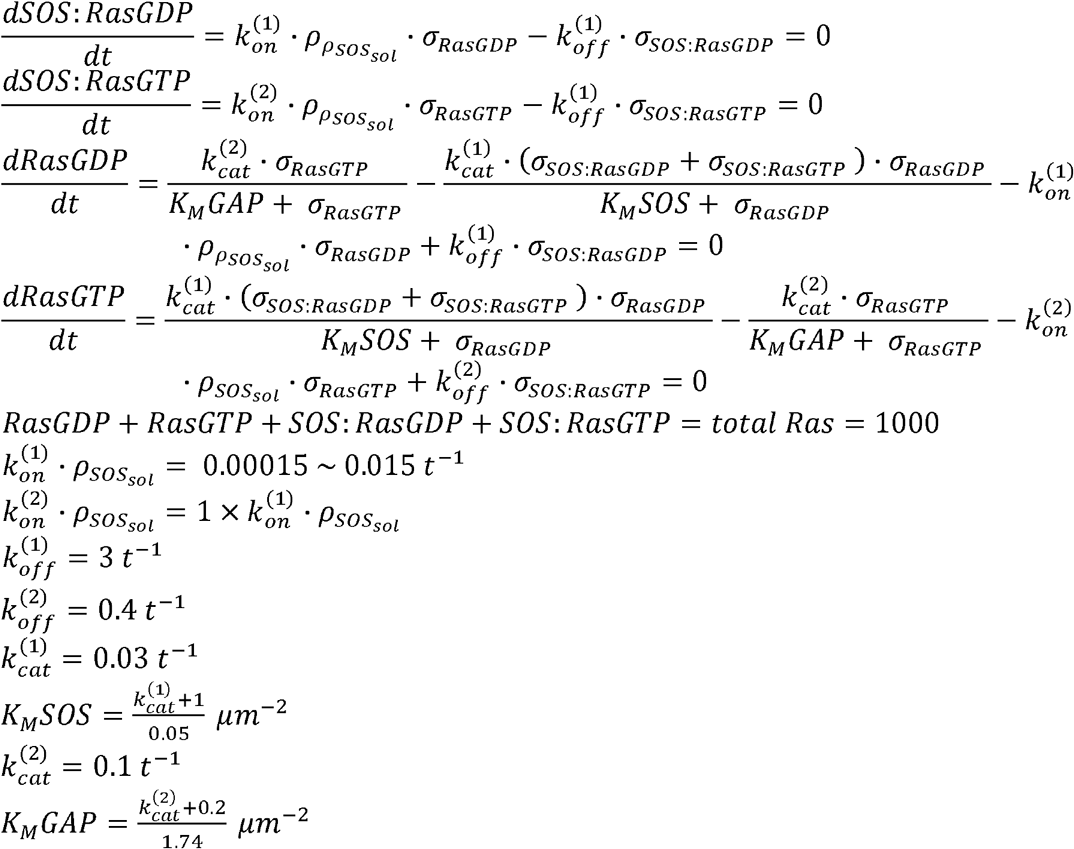

### Statistical Analysis

Non-unimodal conditions were identified based on the Hartigans’ dip test (p < 0.01).

## Acknowledgments

We thank John Kuriyan for providing the plasmids for SOS derivatives and discussions. We thank the members of the Groves Laboratory for helpful discussion and critical feedback on the manuscript.

## Author contributions

Conceptualization: AAL, JTG

Reagent preparation: AAL, SA, JBD, LJNL

Experimental data collection: AAL, HR

Kinetic modeling: AAL, NHK

Data analysis: AAL, NHK, HR

Writing: AAL, JTG

## Data and materials availability

All data are available in the main text or the supplementary materials.

