## Supplementary material for "Bimodality in Ras signaling originates from processivity of the Ras activator SOS without classic kinetic bistability": SI

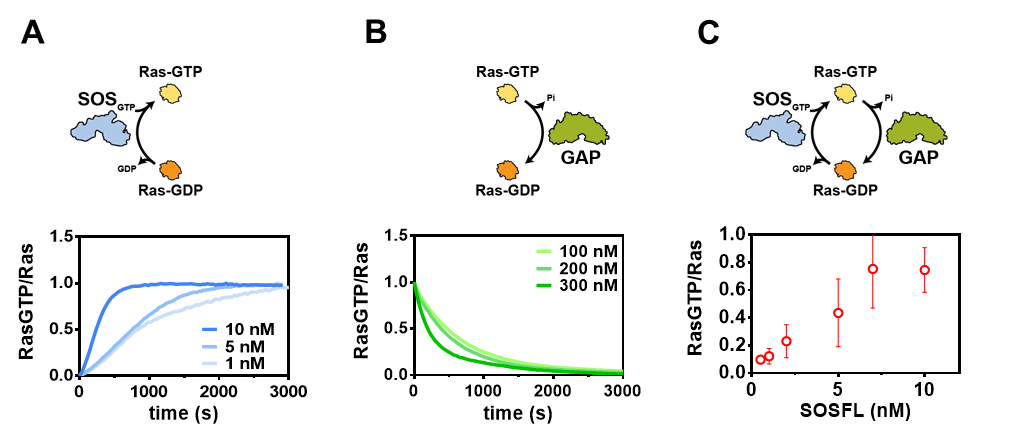

**Fig. S1. Reconstituted Ras activation and deactivation reaction. (A)** Ras-GDP to Ras-GTP nucleotide reaction catalyzed by SOS. **(B)** Ras-GDP to Ras-GTP nucleotide reaction catalyzed by RasGAP. **(C)** Steady state response of Ras activation-deactivation reaction with 200 nM RasGAP and varying concentration of SOS.

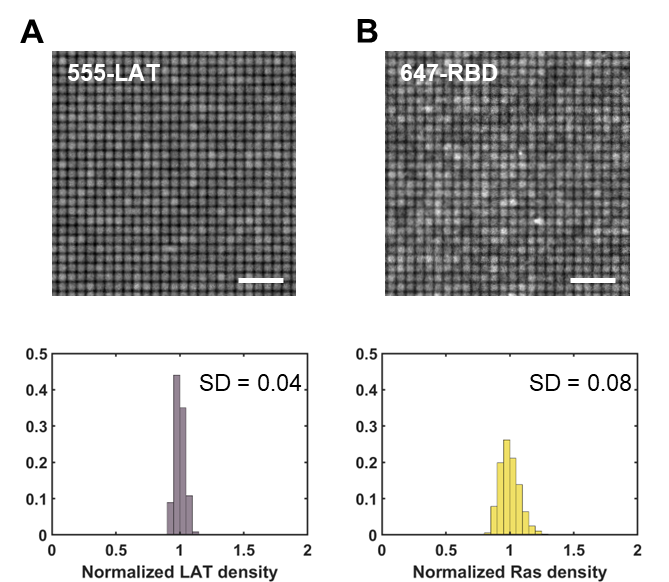

**Fig. S2. Density variation analysis in the micropatterned supported lipid bilayer. (A)** Intensity distribution of 555-LAT across corrals (n>1000). Scale bar 5 μm. **(B)** Intensity distribution of Ras (measured by 647-RBD after Ras was fully activated by SOS to Ras-GTP) across corrals (n>1000). Scale bar 5 μm.

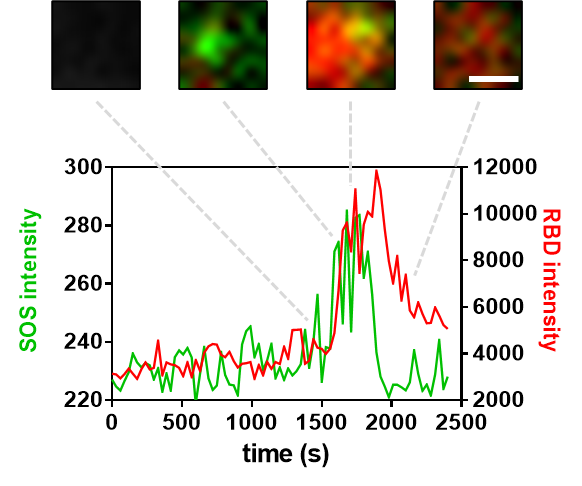

**Fig. S3. Representative example of a burst in Ras activation caused by a single molecule SOS.**. The images and plot are from a single 1 μm × 1 μm corral in a Ras activation-deactivation reaction performed with Alexa-555 labeled SOS. The step increase in the SOS intensity indicates the recruitment of a single SOS molecule to the membrane, as shown in the images above. This is followed by a rapid increase in RBD intensity, indicative of a burst in Ras activation. After SOS detached form the membrane, the Ras activation slowly decreases from the RasGAP activity.

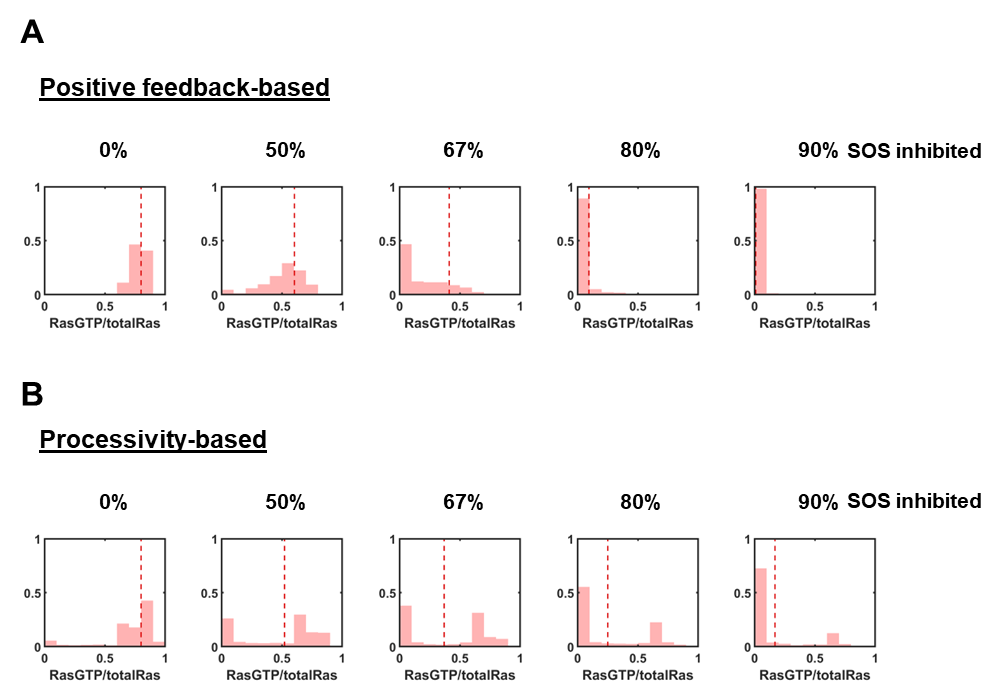

**Fig. S4. Simulation of SOS inhibition. (A)** Positive feedback-based stochastic bistability scenario with SOS inhibition. Distribution of Ras activation state with varying amount of SOS inhibitor in the stochastic simulation (red histogram) plotted with the deterministic steady state solution (dotted red line). At no inhibitor (0% SOS inhibited), Ras activation is 0.7 according to the deterministic steady state solution. At 90% SOS inhibited, both the stochastic simulation and deterministic steady state solution indicates successful inhibition of Ras activation. **(B)** Processivity-based stochastic bistability scenario with SOS inhibition. Distribution of Ras activation state with varying amount of SOS inhibitor in the stochastic simulation (red histogram) plotted with the deterministic steady state solution (dotted red line). At no inhibitor (0% SOS inhibited), Ras activation is 0.7 according to the deterministic steady state solution. At 90% SOS inhibited, while the deterministic steady state solution suggested successful inhibition (Ras activation lowered to 0.169) the stochastic simulation showed a fraction of the reactions still exhibits strong activation way above the mean.

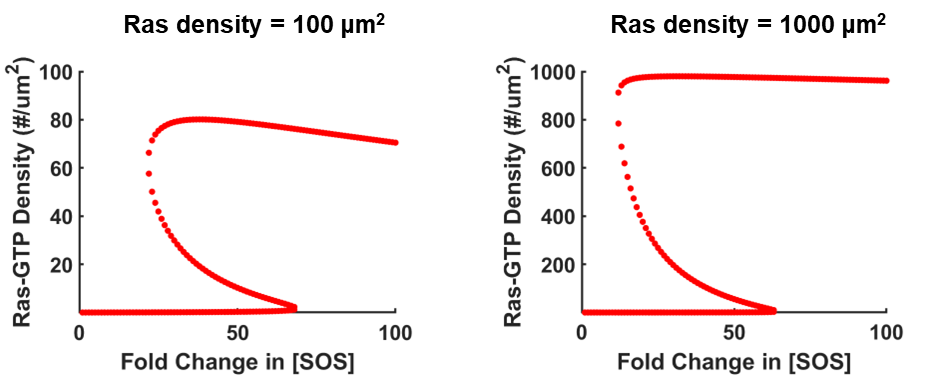

**Fig. S5. Ras activation reaction with kinetic bistability at varying Ras densities.** Using kinetic parameters that leads to kinetic bistability, the deterministic steady state solution can be solved at reaction conditions with either high or low Ras densities. Left: Ras density at 100 μm^2^. Right: Ras density at 1000 μm^2^. The bistable response is robust across different Ras densities.
